# Fast and accurate disulfide bridge detection

**DOI:** 10.1101/2023.11.22.568205

**Authors:** Søren Heissel, Yi He, Andris Jankevics, Yuqi Shi, Henrik Molina, Rosa Viner, Richard A. Scheltema

## Abstract

Recombinant expression of proteins, propelled by therapeutic antibodies, has evolved into a multi-billion-dollar industry. Essential here is quality control assessment of critical attributes such as sequence fidelity, proper folding, and post-translational modifications (PTMs). Errors can lead to diminished bioactivity and, in the context of therapeutic proteins, an elevated risk for immunogenicity. Over the years, many techniques were developed and applied to validate proteins in a standardized and high-throughput fashion. One parameter has, however, so far been challenging to assess. Disulfide bridges, covalent bonds linking two Cysteine residues, assist in the correct folding and stability of proteins and thus have a major influence on their efficacy. Mass spectrometry promises to be an optimal technique to uncover them in a fast and accurate fashion. In this work, we present a unique combination of sample preparation, data acquisition and analysis facilitating the rapid and accurate assessment of disulfide bridges in purified proteins. Through microwave-assisted acid hydrolysis (MAAH), the proteins are digested rapidly and artifact-free into peptides, with a substantial degree of overlap over the sequence. The nonspecific nature of this procedure, however, introduces chemical background which is efficiently removed by integrating ion mobility preceding the mass spectrometric measurement. The nonspecific nature of the digestion step additionally necessitates new developments in data analysis, for which we extended the XlinkX node in Proteome Discoverer (XlinkX/PD) to efficiently process the data and ensure correctness through effective false discovery rate correction. The entire workflow can be completed within one hour, allowing for high-throughput, high-accuracy disulfide mapping.

---

Protein expression describes the intricate processes through which proteins are generated, modified, and regulated in living organisms. Within the wider context of protein research and pharmaceutical applications, these processes are utilized by researchers to direct organisms to produce desired proteins in a process called recombinant expression. Here, plasmids are introduced encoded with the desired protein sequence that, after introduction, are automatically translated into proteins in large amounts. Different organisms can be selected for protein production, each with their own benefits. For example, *E. coli* is the most commonly used organism as it rapidly expands and represents the most cost-effective option. This organism, however, lacks the capabilities to introduce complex post-translational modifications (PTMs) like glycosylation, which can be essential for bioactivity. Other organisms, like Chinese hamster ovary (CHO) cells or insect cells are used in those cases, although these exhibit slower rates of expansion resulting in a higher cost. The advent of recombinant expression has been instrumental to kick-start the biopharmaceutical revolution, which has led to a rapid expansion that is still ongoing. Notably, between the years 2018 and 2022 alone, a total of 180 novel protein products secured regulatory approval[1]. Analysts anticipate further growth due to the mounting incidence of cancer, hereditary disorders, and autoimmune maladies, complemented by the approval of numerous therapeutic interventions that modify the course of these afflictions.

An essential step in protein production is quality control of the product. During this step, the proteins are investigated for multiple parameters to ensure bioactivity and, in the case of biopharmaceuticals, that they cause no immunological issues. Sequence fidelity tests determine whether the proteins have the intended amino acid sequence. Problems that can occur include point-mutations or, in the case of eukaryotic cells, splice variants which can cause loss of activity. Typically, mass spectrometry approaches are used for this step[2]. Folding tests determine whether the secondary and tertiary structure of the protein are correct, the shape of which is essential for bioactivity. Many structural biology techniques are employed here like FRET assays[3] and crystallography or single particle electron microscopy (EM)[4]. Finally, the presence and correct location of the required PTMs is verified. Typically mass spectrometry approaches are used.

Disulfide bridges are covalent links between two cysteine residues existing commonly within the same polypeptide chain (intra-chain links) and less frequently between two polypeptide chains (inter-chain links). This PTM is facilitated in the endoplasmic reticulum by the enzyme protein disulfide isomerase and is amongst the most common PTMs[5]. Present mostly in secreted proteins (a large source of biopharmaceuticals), these links are critical to the protein and incorrect disulfide structures may lead to degradation, loss of function or diseases [6]. Disulfide mapping is one of many characteristics considered as quality attributes that must be thoroughly characterized during protein production. Mismatch of the disulfide bridges (disulfide scrambling) can cause a complex impurity profile and decreased efficacy of the biopharmaceutical which is why official guidelines call for disulfide bridge analysis during production. It is however not trivial to map the often-complex disulfide networks, making disulfide mapping important in both academic and industrial settings.

Several techniques within structural biology have been applied to detect disulfide bridges, such as X-ray crystallography[7] and NMR, however these are costly to execute both in terms of material needed and time spent on the analysis. In recent years, liquid chromatography-mass spectrometry (LC-MS) has emerged as a powerful orthogonal tool for studying disulfide dynamics. In the classic bottom-up proteomics workflow, proteins are reduced and alkylated (effectively disrupting the disulfide bridges) and finally digested into peptides with proteases such as Trypsin. After desalting the peptide mixture is separated by LC and finally analyzed by mass spectrometry where the peptides are fragmented along the peptide backbone in the gas phase[8]. Disulfide bridges can be detected with this technique by either a step-wise reduction and alkylation to determine which cysteine residues are engaged in links [9,10] or by direct detection of disulfide bridged peptides which requires processing the proteins under non-reducing conditions[11].

The unambiguous determination of disulfide bridges through mass spectrometry remains challenging due to several factors such as poor fragmentation properties of linked peptides and limited tools for accurate analysis. Researchers commonly digest the protein(s) using Trypsin, but the alkaline pH optimum makes the disulfide bridges prone to scrambling leading to incorrect results[12]. The aspartic protease Pepsin has a lower cleavage specificity than that of Trypsin, while remaining active at lower pH conditions[13] where disulfide scrambling is less likely to occur and has been used as a more favorable alternative to conventional tryptic digestion in disulfide mapping[14,15]. Proteins however can also be cleaved non-enzymatically by strong acid under high temperatures, either to their individual amino acid components[16] or, by microwave-assisted acid hydrolysis (MAAH), to peptides with optimal properties for mass spectrometric detection[17–19]. MAAH carried out with trifluoroacetic acid (TFA) allows for hydrolysis in less than 10 minutes and is able to provide extensive sequence coverage[18]. As the cleavage specificity is practically random, peptide generation is not reliant on the protein sequence as is the case with conventional proteolytic digestion and a high degree in overlap over the full sequence is typically achieved. MAAH has furthermore shown potential for disulfide mapping[20]. However, the nonspecific cleavage pattern combined with combinatoric analyses make data analysis challenging.

In this work we present a strategy for rapid and confident disulfide mapping using acid-hydrolysis for non-specific backbone cleavage. The highly abundant chemical noise and other singly-charged ions are filtered using FAIMS, which has previously shown great potential in cross-linking MS[21]. Linked peptides are fragmented using EThcD (Supplemental activation to ETD, enhancing dissociation), providing diagnostic ions from the reduced linear peptides along with extensive backbone fragmentation. The data is analyzed using an optimized version of XlinkX[22], capable of FDR-controlled disulfide mapping in mere minutes, allowing for rapid and comprehensive disulfide mapping within 1 hour from intact protein to final result. We validated our workflow by mapping disulfide bridges of 3 standard proteins, chicken lysozyme C, the monoclonal antibody Trastuzumab, and human integrin α-II-b. For all, the disulfide structures are described in literature and represent proteins that are of interest to many different labs studying disulfide mapping.

## Results and discussion

### MAAH digestion properties

As previously reported, MAAH has the capability to break down proteins into peptides suitable for sequencing in under 2 minutes[28]. The drawback is that cleavage is indiscriminate along the entire protein backbone. To enable fragmentation spectrum searches, the cleavage specificity needs to be set as unspecific, or in cases where the search engine does not support this, cleavage after every amino acid (as illustrated in Figure 1A). To control the maximum length of a peptide, the number of allowed missed cleavages is restricted for the latter option to achieve the desired peptide length. By employing the specified search strategy for Lysozyme C samples hydrolyzed for 10 minutes, we achieve nearly 100% sequence coverage (excluding the signal peptide for which no peptides were detected) with peptide ladders across the entire protein backbone. Many of the locations are covered by multiple peptides supported by 100’s of peptide spectrum matches multiplying confidence in the identifications. Coverage is only lost, where the Cysteine residues involved in canonical disulfide bridges are located, suggesting that the disulfide bridges remain intact (Figure 1B). This is not surprising, as the conditions for MAAH are ideal to retain the disulfide bridges due to its short reaction and the acidic environment preventing spontaneous formation of disulfide bridges between free Cysteine residues.

**Figure 1.**
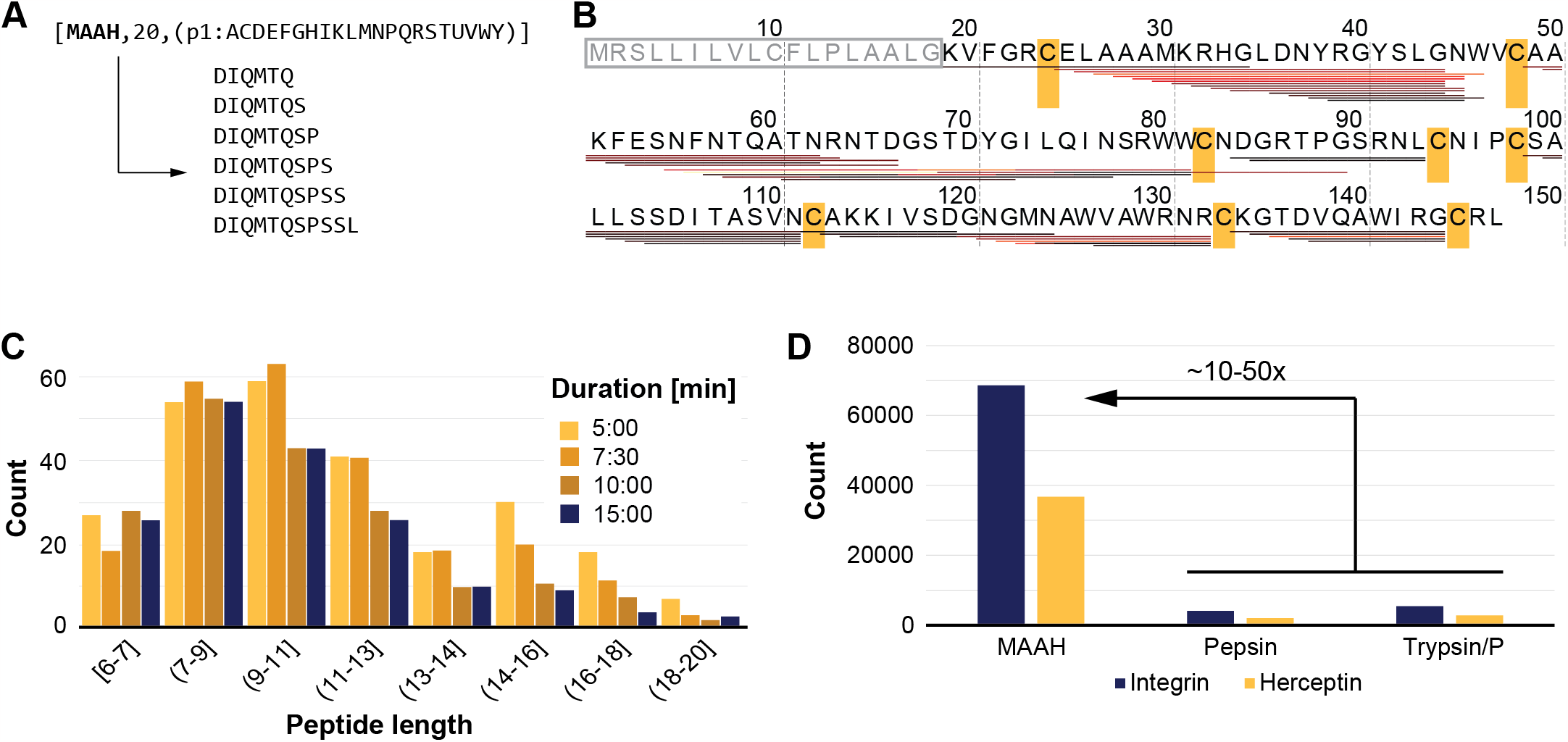
MAAH properties. **(A)** Cleavage settings denoted as [NAME], [MISSED CLEAVAGES], and [SPECIFICITY] in ExPASy PeptideCutter notation. Below example of theoretical peptides. **(B)** Sequence coverage obtained with peptides grouped on first position only. Heat colors denote the number of PSMs at each peptide (ranging from black (n=1) → red (n=100) → yellow (n=250)). Positions of Cys residues involved in canonical disulfide bridges highlighted in yellow. Signal peptide in grey letters inside a grey box. **(C)** Peptide length for different hydrolysis times. **(D)** Effect of protease specificity on the number of theoretical peaks (or search space) generated by the XlinkX/PD search engine.

The presence of the extensive peptide ladders indicates that MAAH indeed cleaves after every possible amino acid residue. Investigating different hydrolysis times (5, 7.5, 10, and 15 minutes) shows that for all times tested, peptides with a length between 6 and 20 amino acids are generated with an average of ∼10 amino acids (Figure 1C).

Interestingly, for the shorter periods a set of larger peptides around 16 amino acids remain, pointing to incomplete digestion. This starts to improve after 10 minutes and eliminated by the 15-minute mark, which we consider the ideal timeframe. Excitingly, an average of 10 amino acids aligns perfectly with the ideal length for peptide search engines. Fragment spectra from peptides that are too short often lack unique assignments to specific peptide identities and peptides that are too long result in low quality fragmentation spectra. All characteristics combined, position MAAH as a potential *idealase* candidate, a digestion strategy dependent not only on specific residues but also on length or size [29]. However, MAAH as a digestion strategy comes at a price. MAAH introduces modifications, which were investigated using an open search of a Trastuzumab MAAH hydrolysate analyzed by LC-MSMS with HCD fragmentation. The modifications were found to consist mainly of deamidation of N and Q, with minor contributions from other modifications, such as dehydration of S and T, oxidation of M, and loss of ammonia, which was found to affect mostly N and Q (Supplementary Figure S2). Only deamidation was included as a variable modification in subsequent searches as it is the most common modification.

Combined with the unspecific cleavage pattern, this leads to a substantial expansion of the theoretical search space, ranging from at least 10 to 50 times its original size (Figure 1D). Such search space expansion leads to unacceptable search times that requires alterations to the search engine.

### Optimal peptide fragmentation

It is by now well-established that under ETD conditions disulfide bridges are more readily reduced than the protein backbone is cleaved[30]. This property is extensively used to improve the sequence coverage in fragmentation spectra in those cases where the disulfide bridges are not reduced during the sample preparation (for example native mass spectrometry experiments). The gas-phase reduction of disulfide bridges results in diagnostic ions that exhibit a specific mass difference of 1.0079 Da for each peptide individually. Attempts have been made to make use of these diagnostic ions to identify disulfide bridged peptides, but these were less than successful and failed to gain traction as the mass difference of 1 Da is less than ideal for shotgun proteomics experiments. Multiple events can lead to the same mass difference, for example incorrect assignment of the monoisotopic mass, deamidation, amino acid substitutions, and others. On the background of this, the reaction pathway is well understood and a complete picture has emerged (Figure 2A) [30,31]. Looking for disulfide bridges peptides in a MAAH digest with classical crosslinking searches uncovered ‘crosslinked’ peptides where the linker is defined as a disulfide bridge (loss of 2 hydrogens) (Figure 2B). In the fragmentation spectra, we indeed find very abundant diagnostic ions. However, with ETD alone,peptide backbone fragmentation events are not favored leading to low sequence coverage for the peptides individually. Integrating an HCD event, through supplemental activation, improves this, leading to high sequence coverage with c-, y-, and z-ions[14].

**Figure 2.**
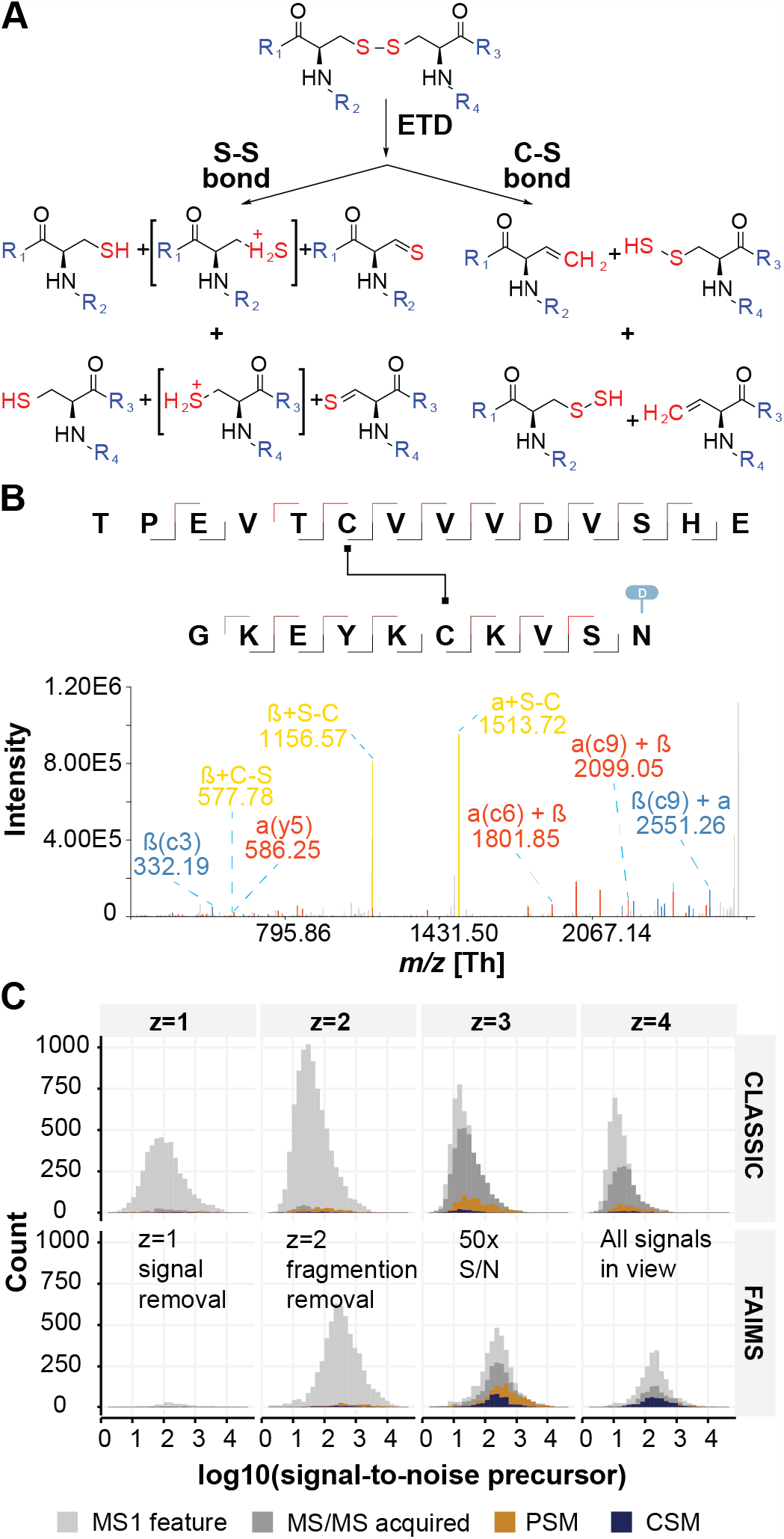
Mass spectrometry optimizations to support disulfide bridge detection. **(A)** ETD driven gas-phase disulfide bridge reduction. **(B)** Example spectrum of disulfide bridged peptides. **(C)** Effect of FAIMS integration into the acquisition investigated as a function of signal-to-noise.

### Ion mobility assisted data acquisition

During the analysis of acid-hydrolyzed proteins we noted that the chromatograms were dominated by highly-abundant singly charged ions. Apart from short, singly charged peptides, these ions are thought to originate from the hydrolysis process itself as they appear in the analyses of several different proteins. The overwhelming intensity of these ions has the potential to mask signals from disulfide-linked peptides via ion-suppression. Based on recent reports, FAIMS has shown great potential in proteomics experiments[32].

A total of 1µg hydrolyzed Trastuzumab was separated across a 60-minute gradient with and without FAIMS applied. Two CV combinations of -50/-60V and -60/-75 were evaluated against a control with no FAIMS. Interestingly, there were no obvious changes to the in the LC-MS profiles (Supplementary Figure S3A), however the scores for the identification increased dramatically (Supplementary Figure S3B). Although the overall intensity in the obtained chromatograms were lower when applying FAIMS, the number of triggered MS2 scans were found to be comparable (15580 without FAIMS and 15054 with FAIMS). The number of observed features with a single positive charge were drastically reduced at both CV combinations, while features from doubly charged species remained numerous (Figure 2C). Interestingly, the S/N of CSMs was increased dramatically for both CV combinations for charge states +3, +4, and +5, which are the common charges observed for disulfide bridged peptides, resulting in a total 5.5-fold increase of CSMs for CV combination -50/-60V and 4.9-fold for -60/-75V over the no-FAIMS control, with an increased number of CSMs associated with every disulfide bridge (Supplementary Figure S3C). Excitingly, this increase brings all signals into view as evident from the transformation of a log normal distribution for the NO FAIMS experiments to a normal distribution for the FAIMS experiments.

Compared to enzymatic digestions with trypsin, MAAH routinely allows for detection of several hundreds (in some cases thousands) of peptides for each protein, meaning each residue is covered by a high number of peptides. The same is true for disulfide bridged peptides, and the high number of repeated CSMs per link were found to be efficient in FDR controlling on the cross-link-level. The increased number of CSMs obtained by adding FAIMS was thereby crucial for detecting false positives and increasing the accuracy of the strategy.

### Search engine optimization

To increase the efficiency of data analysis, and overcome the challenges posed by the unspecific nature of MAAH protein digestion, we integrated an open search module into the XlinkX/PD data analysis environment (see Materials and Methods). Despite working exclusively with purified proteins, the expanded theoretical search space (see MAAH properties) resulted in data analysis times exceeding 24 hours per sample using traditional methods [22,33]. To streamline the data analysis pipeline, we utilize the structure depicted in Figure 3A. The pipeline starts with deisotoping and cleaning of fragmentation spectra extracted by Proteome Discoverer, which already includes accurate precursor *m/z* and charge-state information. Further steps are as follows:

**Figure 3.**
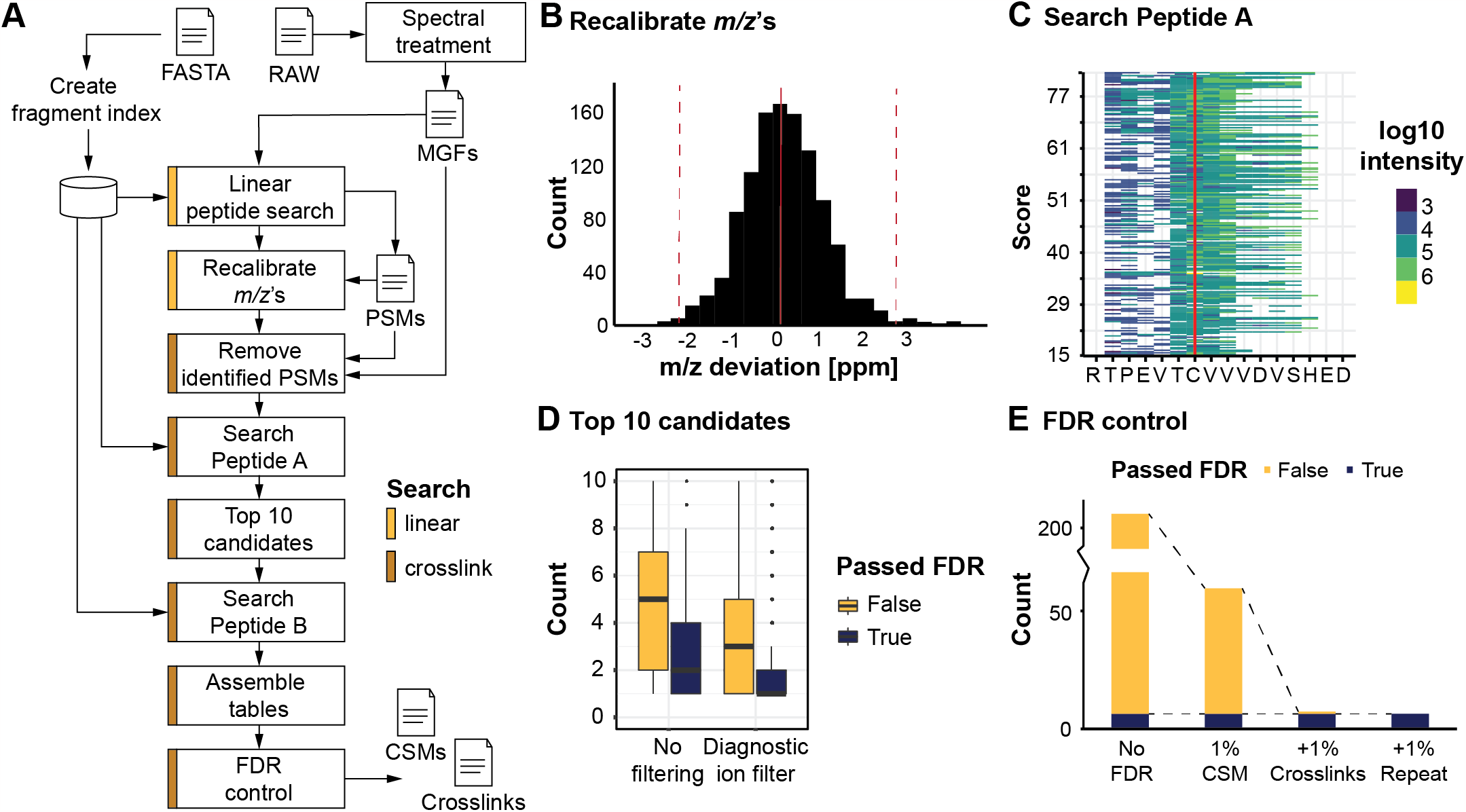
Data processing steps and their results in XlinkX/PD. **(A)** Workflow for processing RAW data with the challenging conditions imposed by MAAH digestion and producing the final Crosslink Spectrum Match (CSM) and Crosslink tables. Linear peptide search in yellow; crosslinked peptide search in brown. **(B)** The precursor *m/z* values resulting from the step “Recalibrate *m/z*’s” (in the RAW file ∼5 ppm off). The boundaries (dotted red line) are estimated with interquartile range fences. **(C)** Sequence coverage with c-ions obtained during the step “Search Peptide A”. Each line is a single fragmentation spectrum, 218 identifications in total. **(D)** The rank of the correct Peptide A after sorting on score differentiated on whether or not including knowledge on the cleavable nature of the crosslink/disulfide. **(E)** Effectivity of the implemented FDR approach.

1. **Cluster Peaks Removal:** Noisy peaks clustering around the real peak, likely artifacts from the Fourier transform, are eliminated;
2. **Precursor Peak Removal:** Peaks larger than the precursor mass minus 2x water are removed to avoid negative impact on the hyper-score used for open search;
3. **Immonium Ions Removal:** These ions are removed based on accurate masses to avoid negative impact on the hyper-score;
4. **Spectrum Filtering:** The spectrum is filtered to retain the TopX of 20 peaks per 100 Da, a standard method to remove noise peaks;
5. **Neutral Loss Peaks Removal:** Non-annotated neutral loss peaks are removed based on mass differences, enhancing the hyper-score accuracy.

Combined, these steps yield clean and interpretable fragmentation spectra.

The second stage focuses on identifying linear peptides from the available fragmentation spectra, as these peptides are easier to identify than crosslinked pairs. After False Discovery Rate (FDR) control at the PSM, Peptide, and Protein level (not shown in Figure 3A), this step provides information for calibrating precursor masses (Supplementary Figure S4A and B). A maximum acceptable mass deviation is estimated after calibration, utilizing interquartile range fences (red dotted lines in Figure 3B). The effect of applying *m/z* calibration is dramatic, as even after stringent FDR control (see next paragraph) false positives remain when it is not applied while only the correct disulfide bridges remain when it is (Supplementary Figure S4C). For the presented data, a development version of XlinkX/PD capable of identifying linear peptides was used. However, similar results can be obtained with regards to mass calibration and removal of confidently identified spectra by using any search engine for linear peptides available in Proteome Discoverer.

In the final stage, crosslinked (or disulfide bridged) peptides were identified. Confidently identified peptide spectra are removed and an open search strategy employed to identify the possible sequence of the first peptide (PeptideA) in the remaining spectra. High sequence coverage for the correct PeptideA was achieved for all spectra supported by intense fragment ions (Figure 3C). This information is then used to create a list of possible peptide identities, sorted based on the hyper-score used in the open search, resulting in the correct peptide located within the top 8 in >95% of the cases. Enforcing the presence of at least one ‘crosslinker’ cleavage product improves the accuracy significantly, with the correct identity in over 95% of the cases located within the top 3. The second peptide (PeptideB) is identified by calculating the mass difference between the precursor and the summed mass of PeptideA and the crosslinker, followed by a classical search approach using a scoring routine (see Materials and Methods). Due to only considering the top 10 PeptideA identities, search time is significantly reduced from excess of 24 hr to less than 5 minutes per raw-file.

### False Discovery Rate correction

In the final stage of the analysis, we compile the complete list of identified crosslinked peptide spectra, along with their corresponding peptide pairs, into a CSM table. However, this table contains, next to the canonical disulfide bridges, numerous false positives (depicted in Figure 3D) that require elimination through a curation process. To achieve this, we employ a multi-stage strategy as outlined by Lenz *et al*[34]. Initially, we identify the peptide identities for both PeptideA and PeptideB using a database containing authentic protein sequences and another with decoy sequences (comprising fully reversed protein sequences, digested into theoretical peptides). Only the top-scoring pair is retained for analysis. If either of the peptides originates from the decoy database, the identification is flagged as decoy. To establish the appropriate score cutoff, the complete list is sorted, and the number of decoy identifications is calculated for each potential cutoff, following the methodology described by Elias *et al*[35]. This step eliminates more than half of the false positives, although the remaining fraction necessitates further curation. The results from the CSM table are then condensed into a crosslinks table, grouping peptide-pairs based on their positions in the protein, including PTMs and missed cleavages. Similar to the CSM table, the crosslinks table is FDR controlled based on the maximum score among all entries for each crosslink. These filtering steps effectively eliminate false positive identifications, which are common due to the nonspecific nature of the digestion and the significant expansion of the search space (up to 50-100 times). Consequently, only canonical disulfide bridges remain, with just one false positive identification.

### Disulfide bridge scrambling

Given that our approach is able to uncover the canonical disulfide bridges, we investigated its ability to detect scrambled disulfide bridges (*i*.*e*., non-biologically relevant disulfide bridges induced by incorrect folding or sample preparation issues). This ability is critical if the workflow is to be used as a QC tool. To ascertain this, we evaluated the degree of disulfide scrambling in a Lysozyme C hydrolysate incubated at pH 8.5 at room temperature (25°C), 37°C and 50°C for 1, 3, and 6 hours; optimal conditions for free Cysteines to react with one another (Figure 4A). As expected, we exclusively identified the four correct bridges in the control sample[36]. At room temperature, however, we already observed three scrambled bridges after 6 hours of incubation. At 37°C and 50°C, after just one hour incubation we observed four and 17 scrambled bridges, respectively. The number of scrambled disulfide bridges rapidly increases with longer incubation times and after 3 hours at 37°C a total of 20 scrambled disulfide bridges were identified, which is comparable to the number identified after just 1 hour at 50°C. We noted that that the intensities of the canonical disulfide bridges significantly dropped after incubation at pH 8.5. Peptides representing non-canonical links increased rapidly in intensity at 37°C and at 50°C. These links had reached intensities comparable to those of the canonical links within just 1 hour (Figure 4B).

**Figure 4.**
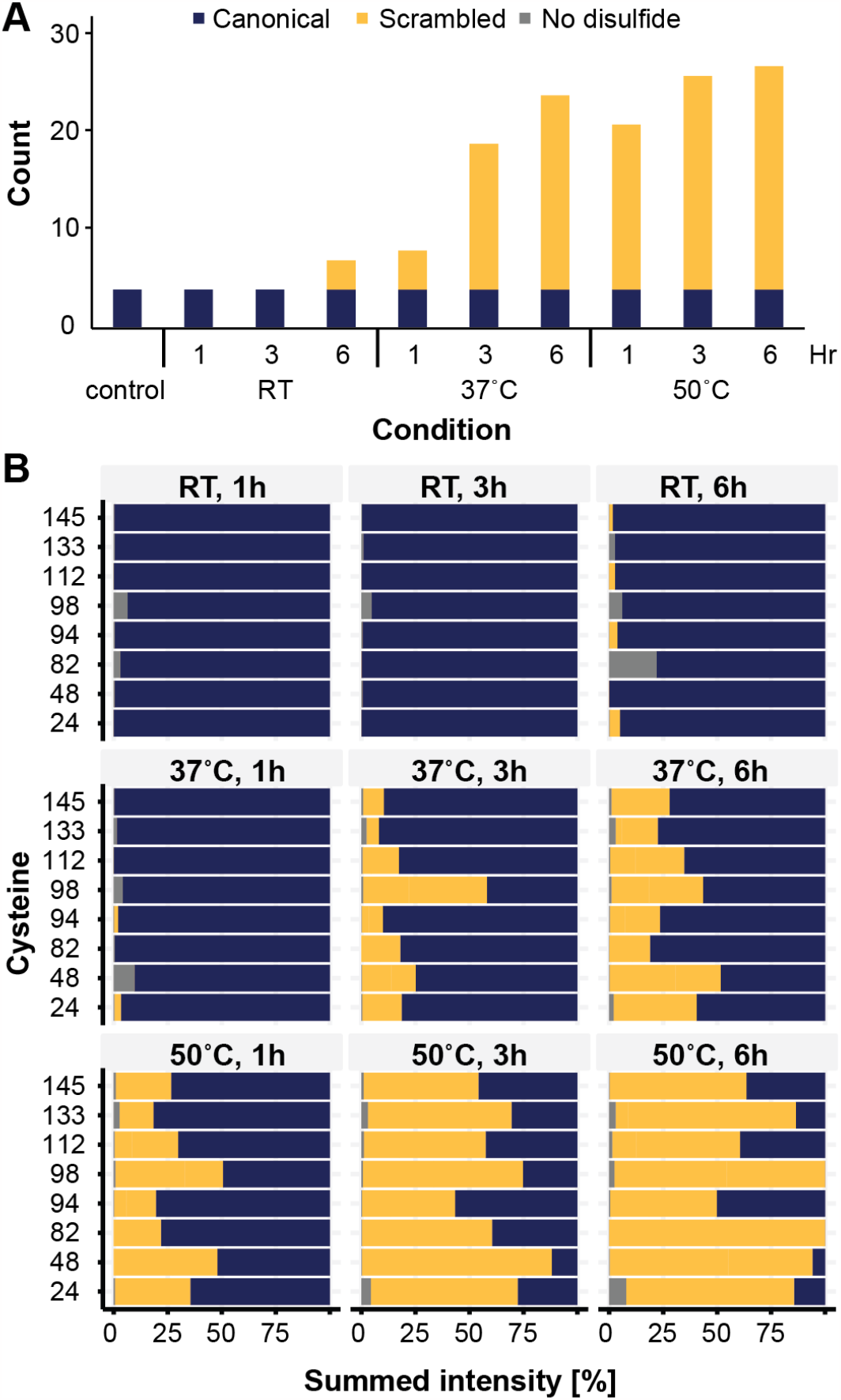
Detection of disulfide bridge scrambling. **(A)** Detected disulfide bridges as a function of the used conditions (top time, bottom temperature). **(B)** Summed intensities of the detected disulfide bridges and linear peptides.

### Disulfide mapping of protein standards

To further highlight the capability of the presented approach, we characterized the disulfide bridges of proteins with a clear clinical relevance and varying complexity in their disulfide landscapes. The first, Trastuzumab, is a monoclonal antibody that is used to treat breast cancer for patients that are shown to be HER2 receptor positive. The second, Human integrin alpha-IIb, is expressed by a wide variety of cell types including T cells (the NKT cells), NK cells, fibroblasts and platelets. Integrins are involved in cell adhesion, participate in cell-surface-mediated signaling and play a critical role in platelet aggregation.

For Trastuzumab, a total of 9 disulfide bridges are described[37] which are indicated in schematic form in Figure 5A. From our analysis, we identify 7 bridges with standard settings. The disulfide bridges at Cys229 and Cys232 are missing, which can be explained by their close spacing, resulting in peptide pairs connected by two disulfide bridges. To account for this, we added a disulfide bridge (*i.e*. H(-2)) as variable modification, resulting in the positive identification of these disulfide bridges in a single peptide pair. The calculated occupancy rates show high occupancy for all disulfide bridges except for one (Figure 5B). The bridge Cys370-Cys428 is in 86% of the cases occupied. It was previously shown that, though removal of this disulfide bridge affects stability of the antibody, it does not structurally change the antibody to the level that intact IgG can still be formed and the antibody remains active[38]. This is well supported by its role in stabilizing two already connected beta sheets (Figure 5C). Hexamerization can be affected potentially modulating complement activation, although previously it was observed that disconnected disulfide bridges can spontaneously reform in the presence of plasma[39].

**Figure 5.**
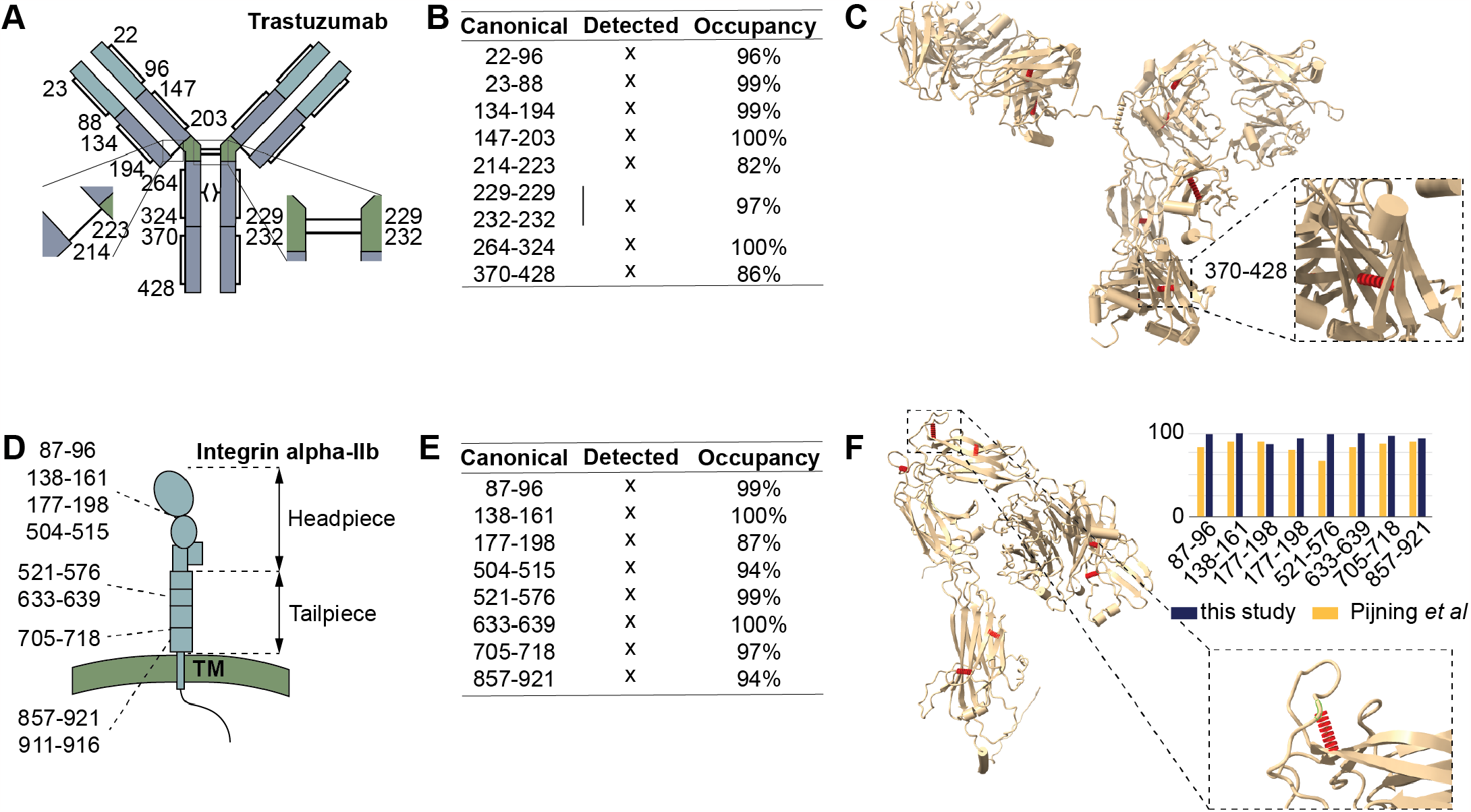
Disulfide mapping of relevant proteins. **A**. Schematic of Trastuzumab with the canonical disulfide bridges indicated. **B**. All canonical disulfide bridges were detected (numbering in uniport, for EU numbering subtract 3) and found to have near 100% occupancy rate, except for the disulfide bridges 214-223 370-428 (middle panel). **C**. Structure of Trastuzumab with the disulfide bridges highlighted (inset: zoom-in on the structure around 370-428). **D**. Schematic of Integrin alpha-IIb with the disulfides indicated in the domains. **E**. All canonical disulfide bridges were detected and found to have high occupancy rates. **F**. Structure of Integrin alpha-IIb with the disulfide bridges highlighted. Top right, comparison of the occupancy rates of this study and those obtained by Pijning *et al*[40]. Bottom right, zoom-in on structure around Cys504-Cys515, which showed the lowest occupancy rate).

For Integrin alpha-IIb, also 9 disulfide bridges are described[40] that are indicated in schematic form in Figure 5D. With standard settings eight were successfully detected, all at high occupancy rates (Figure 5E). The disulfide bridge Cys911-Cys916 was not found in our analysis, for which previous reports also do not provide evidence of existence[40]. The lowest occupancy is detected for Cys504-Cys512, which maintains a small loop (Figure 5F). To verify whether our observation is correct we correlated our occupancy rates to those previously reported by Pijning et al. Also this study reports Cys504-Cys512 with the lowest occupancy, while the rest of the disulfide bridges correlate very well on occupancy rates.

### Analysis time reduction

Sample throughput is major concern in many proteomics laboratories, both industrial and academic. Given the significant speed increase in data analysis and the extreme repetition rates of identifications, we reasoned that the data acquisition time can be reduced as well. To this end, we evaluated the capabilities of this approach using a 20-minute and a 30-minute gradient rather than 60 minutes, which would allow for disulfide mapping on a much shorter time scale. Considering the drastic increase in data quality attributed to FAIMS we hypothesized that a shorter gradient would still provide extensive linkage information. Using Lysozyme C we were able to cover all 4 disulfide bridges using both the 20-minute, 30-minute, and 60-minute gradient with 79, 135, and 217 CSMs respectively. These results demonstrate that we can go from intact protein to complete disulfide mapping results within a single hour.

## Conclusions and Discussion

In this paper, we present an efficient and precise sample preparation, data acquisition and analysis approach for the accurate detection of disulfide bridges. In earlier studies, researchers attempted to identify disulfide bridges through indirect means like differential alkylation or direct detection as ‘crosslinked’ peptides[41–48]. Typically, these methods involved analyzing samples under unfavorable conditions. For instance, they employed traditional enzymatic reactions where digestion occurred in a slightly alkaline pH setting. While this approach allowed the utilization of existing tools tailored for proteomics workflows, it had a drawback: it left proteins susceptible to rearrangement of their disulfide bridges, forming non-biologically relevant connections between Cysteines. To overcome this, several studies report reduced scrambling when performing digestions at pH 5-6, by applying proteases with a lower pH optimum such as pepsin[14,15], or by the addition of an oxidizing agent[49]. Most enzymes used in proteomics experiments however require higher temperatures and longer incubation times to ensure complete digestion and, so far, these did not gain traction for the detection of disulfide bridges. In contrast, the MAAH that is central to our approach efficiently hydrolyzes proteins in highly acidic conditions, preventing such scrambling. MAAH however produces a high abundant background which must be removed to keep required high dynamic range for disulfide identification. Adding FAIMS as a gas filtering device improves the number of identified CSMs 5 folds (Figure 2) and enables identification of all S-S links in the systems we investigated. EThcD has previously shown great potential for disulfide analysis[14] and as this fragmentation strategy produces not only backbone fragments but also intense ions from the intact linear peptides, the spectra were rich in information.

The required search engine optimizations were implemented in the XlinkX node in PD v. 3.1 sp1. This data analysis pipeline has for the last 7 years gone through continuous development cycles and will continue to do so. For the near future, we are extending its functionality to encompass a broader set of options. One of these will be the extension of XlinkX/PD for the extraction of higher-order (more than two) crosslinked peptides. Currently, XlinkX/PD is restricted to two peptides (or di-peptide) crosslinked together. For disulfide bridged peptides more peptides need to be supported. A famous example is Insulin, where the disulfide bridges intertwine around closely spaced positions. For the current implementation a workaround enables their detection.

Our comprehensive protocol not only accelerates the sample preparation, data acquisition, and data analysis but also guarantees precise and dependable identification of crosslinked peptides in intricate samples. This makes it an invaluable tool for high-throughput quality control applications, which is an increasingly important part of the biopharmaceutical pipeline. At a current estimated market share of 450 billion dollars annually, which is anticipated to exceed 1 trillion dollars by 2030, our protocol can become a key component in the biopharmaceutical pipeline. Disulfide mapping is also a critical attribute in detailed protein structure characterization. As currently most knowledge about disulfide bridges comes from much more complicated classical structural biology experiments like X-ray crystallography or EM, complementary information provided by mass spectrometric analyses can be beneficial. Most proteins analyzed by the classical techniques are overexpressed recombinant and very often alkylated (alkylation improves the resolution for these imaging techniques) proteins that can result in errors of disulfide mapping due to scrambling or alkylation. Mass spectrometry can provide disulfide mapping for more complex and endogenous samples.

## Methods and Materials

### Chemicals and Reagents

Chicken lysozyme C (L-7651) was purchased from Sigma Aldrich. Trastuzumab was generously donated by a pharmaceutical company and human integrin α-IIb was purified from human platelets. Trifluoroacetic acid (TFA), LC-MS grade was purchased from Thermo Scientific.

### Microwave-assisted acid hydrolysis

20 µg of dry protein sample was dissolved in 40 µL 25% TFA in a 1.5mL low-binding polypropylene vial (Eppendorf). The vial was sealed with a micro tube cap lock (Scientific Specialties) and additionally secured with tape before placing it in a bubble rack, which was placed in a 1000mL beaker containing 200mL demineralized water. The beaker was positioned off-center in a household microwave oven (General Electrics, model PEM31DMWW) and hydrolysis was performed using the standard setting at 800W for 15 minutes. Hydrolyzed protein was quickly dried by vacuum centrifugation and dissolved in LC-MS loading solvent (0.1% TFA) prior to injection on the mass spectrometer.

### Assessing disulfide scrambling

Chicken lysozyme C was acid hydrolyzed and purified by in-house constructed zip-tips. Aliquots were dissolved in 50mM TEAB pH 8.5 and incubated at room temperature, 37°C, and 50°C. Time points were taken after 1 hour, 3 hours, and 6 hours and immediately acidified. All time point aliquots were compared to a control sample which had kept at low-pH conditions.

### LC-MS data acquisition

Samples were separated by reverse phase-HPLC using a Thermo Scientific™ Vanquish™ Neo system connected to an EASY-Spray™ PepMap™ RSLC C18 column (0.075 mm x 250 mm, 2 µm particle size, 100 Å pore size (Thermo Fisher Scientific)) at 250 nL/min flow rate. The hydrolyzed samples were analyzed on the Orbitrap Eclipse™ Tribrid™ mass spectrometer coupled with FAIMS Pro Duo interface. Reverse phase separation was accomplished using a 20-, 30- or 60-min separation gradient (plus 10 – 20 min equilibration phase) of 4 – 40% solvent B (A: 0.1% FA; B: 80% ACN, 0.1% FA). FAIMS was set at standard resolution with 3.9 L/min total carrier gas flow and with 2 CV (-50,-60) method. Samples were analyzed using an EThcD-MS2 acquisition strategy with 20% SA collision energy. MS1 and MS2 scans were acquired in the Orbitrap with a respective mass resolution of 120,000 and 60,000. MS1 scan range was set to m/z 375 – 1400, standard AGC target, 246 ms maximum injection time and 60 s dynamic exclusion. MS2 scans in data dependent acquisition mode (top speed 1.5 sec/cv) were set to an AGC target of 2e5, 118 ms max injection time, isolation window 1.6 m/z. Only precursors at charged states +3 to +8 were subjected to MS2.

### Implementation of an open search engine

We implemented an open search engine modelled after the approach described for MSFragger[23], which itself was based on prior work. To speed up fragmentation spectrum annotation, we also make use of a fragment index (Supplementary Figure S1A), where each theoretical fragment is maintained in a sorted list that is binned on 0.01Da intervals. This setup allows for almost instantaneous extraction of peptides matching a particular fragment. All fragments in a spectrum can then be annotated with peptide identities and a list constructed on the most likely peptide identities. To tie everything together additional meta-data is need about the protein sequences, modifications and the total peptide mass (Supplementary Figure S1B). For the initial identification we utilize the hyper-score calculation integrated in Sequest[24]. Based on this score we assemble a top 10 list of the best peptide identities and re-score this with scoring routine adapted from Olsen and Mann[25] to obtain the final peptide identity.

To investigate the performance of the engine, we compared the results from a HeLa cell tryptic digest to those obtained with Mascot or Sequest, frequently used peptide search engines in Proteome Discoverer. We identified approximately 75% of the spectra identified by Mascot along with a very low percentage of additional identifications uniquely identified by our search engine (Supplementary Figure S1C). The remaining fraction of Mascot-identified fragmentation spectra were also correctly identified by our engine but discarded during the FDR control step due to low quality. We conclude therefore that our linear search engine works properly.

### Data analysis

All data analysis, unless otherwise stated, was performed in Proteome Discoverer v. 3.1 SP1. Shortly, the processing workflow consists of the nodes (1) ‘Spectrum files’ to connect the workflow to the raw files. (2) “Spectrum Selector” to extract the fragmentation spectra with accurate precursor *m/z* and charge states. Standard settings applied. (3) Linear peptide search engine, for which we used a development version of the open search linear peptide search engine described in the previous paragraph. This can however be replaced with another engine like Sequest HT or Mascot (see Material and Methods). As FASTA file we used one containing the protein under investigation, that is *in silico* digested with a protease defined as Figure 1A with a minimal peptide length of six and 20 miss cleaved sites allowed. No fixed modification was set. Methionine oxidation, protein N-term acetylation and asparagine & glutamine deamidation were set as dynamic modifications. (4) “Target Decoy PSM Validator” was used to control for false positives at 1% (5) “Spectrum Confidence Filter” was then used to remove all spectra that were confidently identified with the linear peptide search. Setting ‘Worse Than High’. (6) Finally, the XlinkX/PD nodes “XlinkX/PD Detect”, “XlinkX/PD Search”, and “XlinkX/PD Validate” were connected. To define the disulfide bridge, we set up a cleavable crosslinker as H(-2). The ETD cleavage products (or diagnostic ions) are defined as follows: S-S symmetrical as PeptideA=“H(1)” and PeptideB=“H(1)”, C-S asymmetrical PeptideA=“H(2)” and PeptideB=““, S-C asymmetrical full PeptideA=““and PeptideB=“H(1) S(1)”. As ‘Acquisition strategy’ open search was used. The same protein database, and fixed and variable modifications were used as defined in the linear peptide search.

All protein structures were visualized with ChimeraX[26] and the disulfide bridges mapped with the plugin XMAS[27].

## Acknowledgements

We thank Dr. Lu Wang of the Allen and Frances Adler Laboratory of Blood and Vascular Biology for his generous donation of the purified integrin αIIbβ3. RAS acknowledges that this work is part of the research program € TA with project number 741.018.201, financed by the Dutch Research Council (NWO). RAS further acknowledges funding through the European Union Horizon 2020 program INFRAIA project Epic-XS (Project 823839).

## Data availability

Appropriate raw-files and search results have been submitted to ProteomeXchange and can be accessed through accession: PXD046855. For potential review of the data, please use username: “ “and password: “DhS5mphm”. The updated XlinkX node for Proteome Discoverer 3.1 will be made available through a hot fix.

## Contributions

SH, HM, RV, and RAS designed the study. SH, YH, YS and RV performed the experiments. RAS and AJ developed the data analysis toolchain and analyzed the data. SH, RV and RAS wrote the manuscript with help from all authors.

---

**Supplementary Figure S1.**
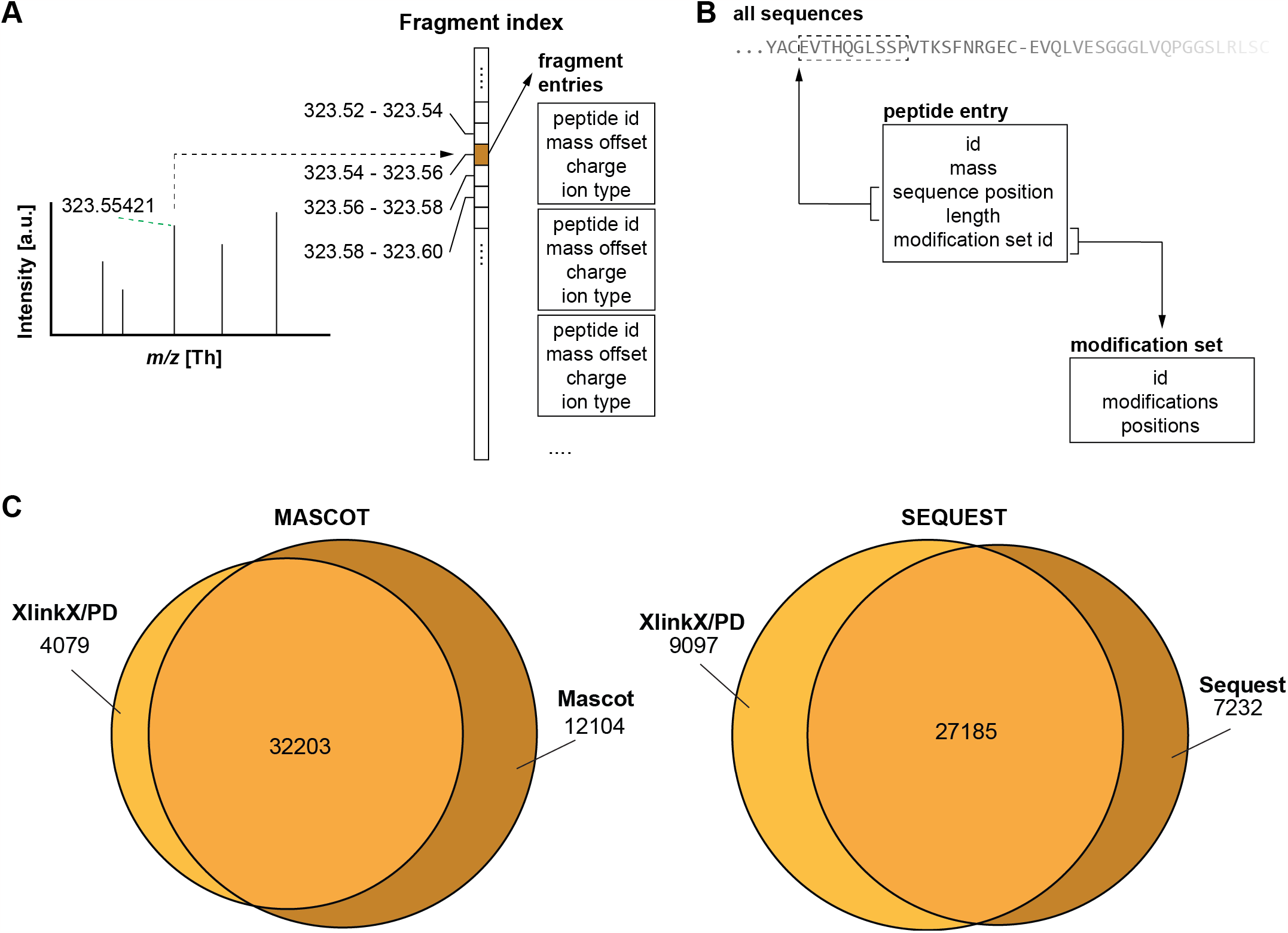
Open search implementation. **(A)** The fragment index implementation initially digests all protein sequences and for each peptide generates the theoretical fragments that are stored in a list sorted on fragment mass. For each peak in the fragmentation spectrum, associated peptides can almost instantaneously be recalled. In our implementation each peptide entry is restricted to 16 bytes, meaning that for 5 million peptides only 80 Mb is used. (**B)** For each peptide meta data is maintained that allows for construction of the sequence and the modification profile. Additionally, an extra list is maintained sorted on mass that allows for a classical approach for searching based on the precursor mass. **C**. When using the search engine in linear peptide search mode allows for comparison to existing search engines. Examination of the overlap of the identified peptide shows good overlap. In cases where the other search engines identify extra peptides in close to all cases our open search detects the same peptide identity but removes them due to quality concerns.

**Supplemental Figure S2.**
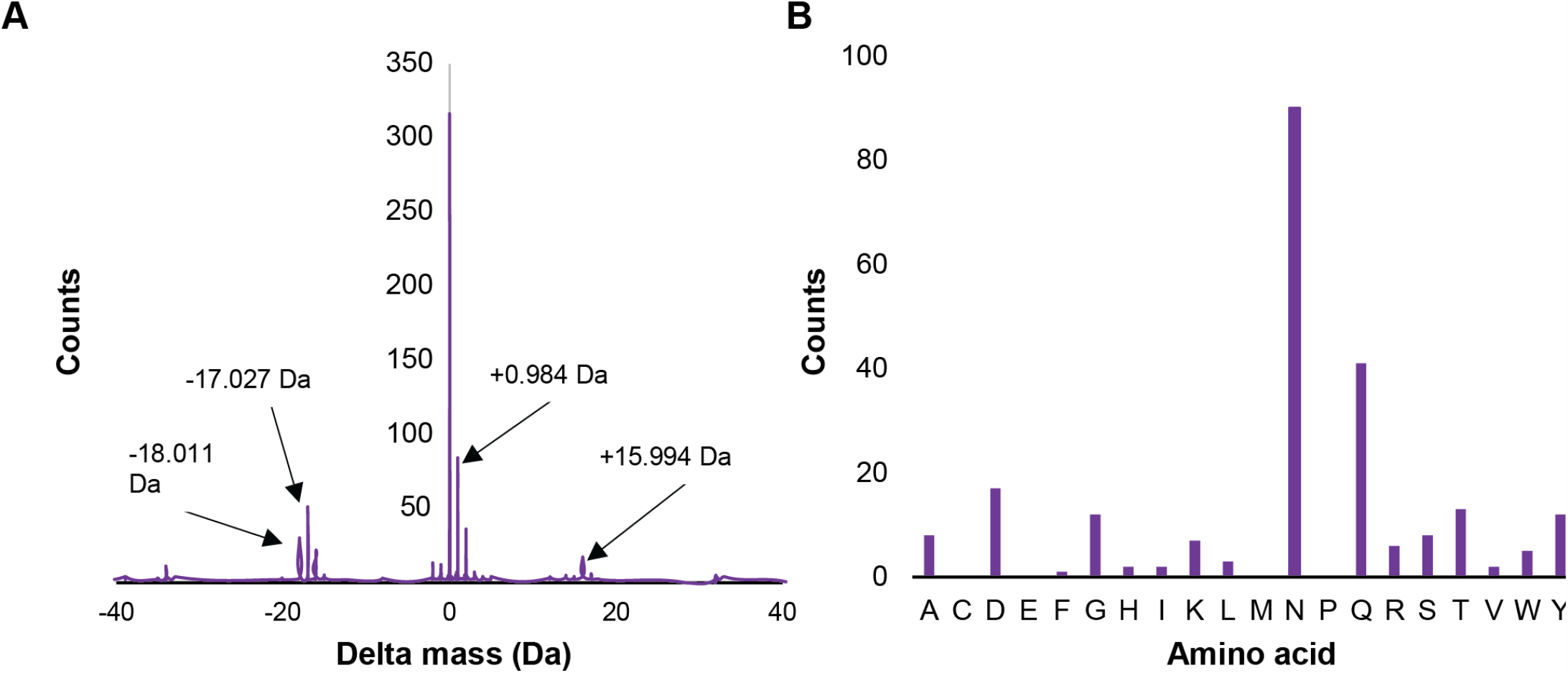
Modifications associated with MAAH. **(A)** Delta masses binned in 0.01 Da intervals plotted against the number of observations. The results were generated from MAAH-hydrolyzed Trastuzumab fragmented with HCD and searched by MSFragger in open search mode. **(B)** Number of -17 Da modifications for each amino acid based on PSMs. Data is produced from MAAH-hydrolyzed Trastuzumab fragmented with EThcD and searched by SEQUEST HT with loss of ammonia (-17.027 Da) as a variable modification potentially inhabiting all residues.

**Supplementary Figure S3.**
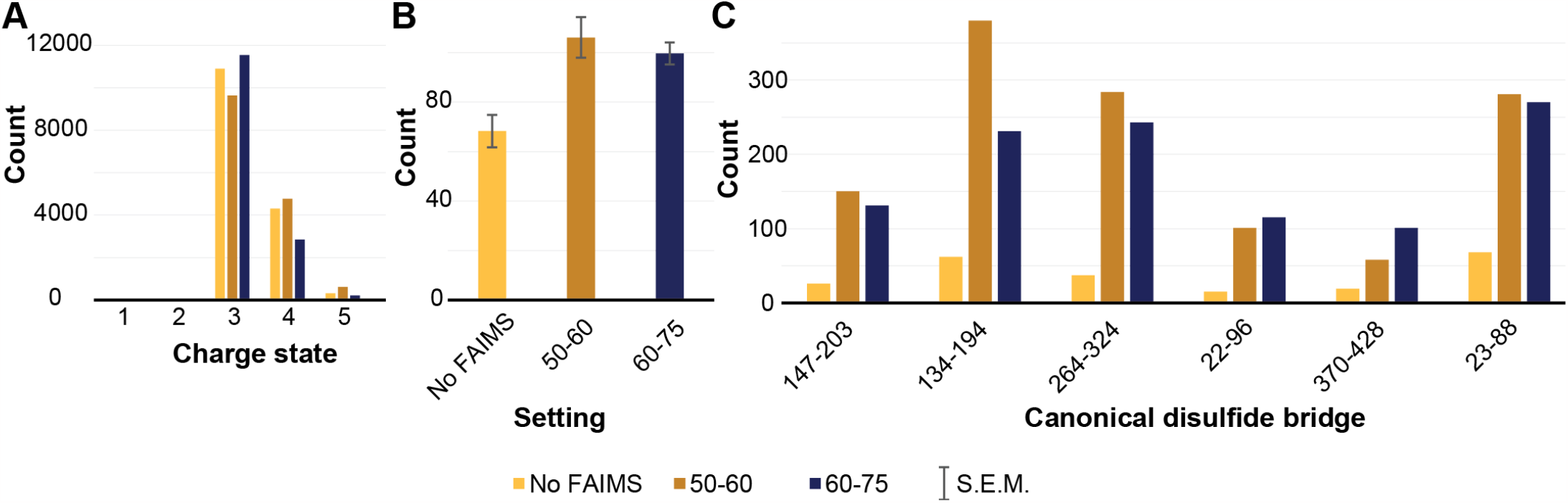
Further details on the effect of FAIMS integration in the data acquisition pipeline. **(A)** Comparing the for fragmentation selected precursor charge states shows no major effect in the selection of the precursor (note, the charge state is that what during the acquisition was determined to be the charge state). **(B)** The scores assigned to the identifications significantly increases between NO FAIMS and FAIMS turned on. This is linked to the increase in signal-to-noise of the precursors as detailed in Figure 2C. **(C)** The improved signal-to-noise of the precursors brings many more precursors in range for successful sequencing, resulting in a significant increase in the number of identified fragmentation spectra per unique disulfide bridge (denoted on the x-axis as the protein positions).

**Supplementary Figure S4.**
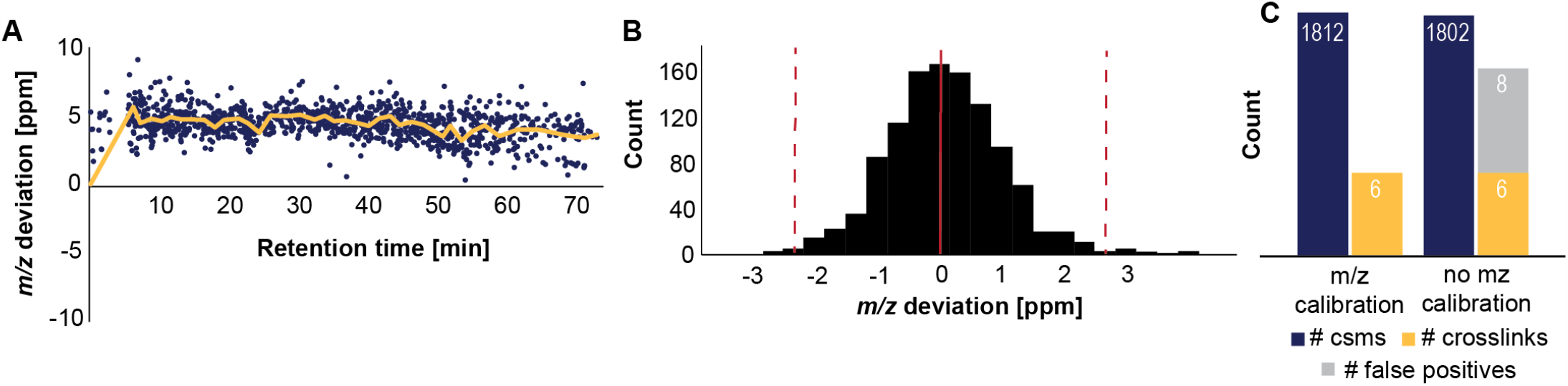
Retention time alignment from peptide identifications. **A**. Examining the difference between the theoretical and the measured precursor monoisotopic mass reveals the *m/z* drift of the instrument. From the point cloud spline can be fit that describes the average *m/z* drift over the retention time. The spline is used to correct the precursor masses of all fragmentation spectra prior to the crosslinked peptide identification step. **B**. After alignment of the precursor masses, the full population adheres to a normal distrubution from which the edges can be estimated in a data driven fashion. We utilize interquartile range fences, where the interquartile range multiplied by a factor of 3 is added (right hand fence) and subtracted (left hand fence) to quartile3. **(C)** When the precursor *m/z* is not applied, false positives can arise even after the stringent FDR controls applied during the data analysis.

